# Narrow-beam geometry improves the efficiency of cryo-EM

**DOI:** 10.64898/2026.07.06.736854

**Authors:** S. Matinyan, P. Filipcik, E. van Genderen, J.P. Abrahams

## Abstract

Cryo-electron microscopy (cryo-EM) of biological specimens is limited by radiation damage and a low signal-to-noise ratio (SNR). Here, we show that reducing the illuminated area substantially slows the observed diffraction decay in protein microcrystals. We further show that narrow parallel-beam electron diffraction from thin non-crystalline biological specimens provides substantially higher reciprocal-space SNR than conventional cryo-EM imaging. We developed a multimodal scanning workflow, 4D-para-STEM, that records narrow-beam diffraction patterns together with corresponding images. Using viruses, peptide assemblies, and microtubules, we demonstrate interpretable diffraction signals from both crystalline and non-crystalline biological specimens. Together, these results show that narrow parallel-beam scanning reduces observed radiation damage and improves the SNR in cryo-EM.

## 1. Introduction

Cryo-electron microscopy (cryo-EM) has transformed structural biology by enabling structure determination of biological macromolecules in near-native states. However, because biological specimens are highly sensitive to electron irradiation, cryo-EM data must be acquired under extreme low-dose conditions [1,2]. Despite major advances in detectors, motion correction, and machine-learning-based denoising, data quality remains dose-limited, with image SNR often remaining in the range of 0.01–0.1 [3,4]. Therefore, even modest improvements in the SNR can reduce the required particle numbers, improve the refinement accuracy, and extend the range of accessible macromolecular targets.

Electron diffraction using a parallel beam can, in principle, be more dose-efficient than conventional imaging [5]. Such diffraction measurements are not attenuated by the contrast transfer function (CTF) and are less sensitive to beam-induced motion blurring and coherence losses associated with inelastic scattering. However, for non-crystalline biological specimens, reciprocal-space SNR depends critically on the illuminated volume: when the beam substantially exceeds the dimensions of the target particle, scattering from the surrounding solvent obscures the weak molecular speckle signal. Narrow parallel-beam illumination minimizes the solvent background by confining the illumination to dimensions comparable to the target structure. Despite this potential advantage, narrow-beam diffraction of non-crystalline biological samples remains comparatively unexplored [6,7].

The importance of beam geometry in reducing radiation-induced information loss was recognized early in cryo-EM. Henderson and Glaeser identified beam-induced specimen motion as a dominant factor limiting high-resolution image contrast in radiation-sensitive materials. Downing and Glaeser (1986) and Bullough and Henderson (1987) subsequently developed spot-scan imaging approaches in which parallel illumination was confined to small regions scanned across the specimen [8,9]. These studies demonstrated substantial improvements in high-resolution contrast and information transfer, in some cases by factors of 3–5, through suppression of beam-induced motion and charging effects [9]. Despite these observations, narrow parallel beam geometry has remained largely underexplored for improving the SNR.

Recent low-dose 4D-STEM and cryo-ptychography approaches explored scanning-based acquisition geometries using narrow *convergent* electron beams in biological samples [10]. In ptychographic reconstruction, structural information is recovered through interference with the undiffracted beam and phase-sensitive image formation. Here, we revisit and extend the concept of using narrow *parallel* beam cryo-EM. We combine diffraction and imaging measurements from identical specimen locations in a multimodal scanning workflow (4D-para-STEM). Using protein microcrystals, viruses, and peptide assemblies, we show that a small beam slows observed radiation damage and enables higher reciprocal-space SNR at lower dose than conventional cryo-EM imaging. Together, these results show that beam size and multimodal acquisition can improve information efficiency in cryo-EM.

## 2. Materials and Methods

### 2.1. Microscope configuration and narrow parallel-beam alignment

Experiments were performed using a JEOL F200 transmission electron microscope in the μμDIFF STEM regime operated at 200 keV and equipped with a Schottky field emission gun, a CEOS CEFID energy filter, a Tietz Scan Generator, a Gatan K2 detector, and an ASI Cheetah M3 hybrid pixel detector. Scanning and detector synchronization were controlled using the Panta Rhei software platform (CEOS GmbH).

Narrow parallel-beam illumination was established using a modified Köhler illumination configuration. Initial parallel beam alignment was performed in standard TEM mode and corresponding lens excitation values were transferred to the STEM mode. Beam parallelism at the specimen plane was optimized by minimizing direct-beam diameter and minimizing beam-size variation during objective lens wobbling. Beam diameters were controlled primarily through condenser aperture selection and third condenser lens (CL3) excitation. The smallest routinely achieved parallel beam diameter was approximately 40 nm.

Separate intermediate lens configurations were calibrated for diffraction mode and for image acquisition at multiple magnifications. Independent scan–descan compensation settings were made for diffraction and imaging conditions to maintain beam centering across the full scan area.

Additional alignment procedures are provided in the Supplementary Materials.

### 2.2. Diffraction and imaging data acquisition

For diffraction acquisition, the detector was configured conjugate to the diffraction plane in STEM mode. Survey scans were initially recorded using virtual dark-field contrast to identify suitable specimen regions. Subsequently, 4D diffraction datasets were acquired as n × n scan grids, yielding n^2^ diffraction patterns per dataset. Typical scans consisted of 100 × 100 positions, with diffraction frames recorded at 514 × 514 pixels.

Imaging datasets were acquired sequentially from identical scan regions using image-plane detector configurations while maintaining constant condenser lens (CL) settings to preserve scan registration between the diffraction and imaging modes. Scan/descan alignment was calibrated independently for diffraction and imaging acquisitions.

For stationary diffraction measurements, diffraction frames were acquired using exposure times of 10 ms at doses of approximately 0.4 e^-^/Å^2^ per frame and were summed computationally, where indicated.

### 2.3. Reciprocal-space SNR estimation and Bragg intensity analysis

Bragg peak intensities and reciprocal-space signal-to-noise ratios (SNRs) were quantified using a custom reciprocal-space analysis framework (Bragg_snr) developed for weak biological nanobeam diffraction data. Unlike conventional crystallographic spot-finding approaches, this framework does not require prior indexing, rotational data collection, or globally consistent lattice geometry, making it suitable for still diffraction frames, partially ordered specimens, multiple lattices, and continuous layer-line diffraction from helical assemblies. Implementation details and mathematical definitions are provided in the Supplementary Materials.

### 2.4. Radiation damage measurements

Radiation damage measurements were performed using lysosome protein microcrystals under parallel-beam illumination conditions. Condenser lens apertures (CLA) of 30, 10, and 5 µm produced calibrated beam diameters of approximately 480, 160, and 80 nm, respectively.

For each beam condition, diffraction patterns from 15–20 crystals of comparable thicknesses were recorded as a function of the accumulated electron dose until the extinction of the Bragg reflections.

### 2.5. Reciprocal-space SNR comparison

Reciprocal-space SNR comparisons between diffraction and imaging data were performed using experimental diffraction patterns and Fourier-transformed cryo-EM images recorded from comparable specimens and spatial-frequency ranges.

Cryo-EM imaging data were acquired on a Titan Krios microscope operated at 300 keV with a nominal pixel size of 0.7384 Å/pixel and a total accumulated dose of 30 e^-^/Å^2^. Images were motion corrected prior to Fourier transformation. Fourier amplitudes were squared to obtain reciprocal-space intensity distributions directly comparable to diffraction measurements. Imaging data were low-pass filtered to match the spatial-frequency range of the diffraction measurements.

Diffraction data were recorded at doses of approximately 4 e^-^/Å^2^. Reciprocal-space SNR values were evaluated from radial line profiles across corresponding layer-line regions.

### 2.6. Virtual dark-field reconstruction and image stitching

Virtual Dark-Field (vDF) images were reconstructed from 4D diffraction datasets by integrating annular diffraction intensities around the direct beam for each scan position. These reconstructions were used to visualize scanned regions and identify positions containing biological signal (Figure S2).

Image patches acquired during scanning were stitched computationally to reconstruct larger fields of view. Image centers were initially estimated using circular Hough transforms. Frames were subsequently aligned using normalized cross-correlation and phase-correlation refinement. Overlapping regions were blended using gradient feathering (Figure S3).

### 2.7. Diffraction-frame decomposition

Diffraction patterns were decomposed into isotropic, centrosymmetric (Friedel-symmetric), asymmetric, and residual noise components to improve extraction of weak biological diffraction signal from diffuse solvent background and localized crystalline diffraction. The procedure exploits radial isotropy of solvent scattering and Friedel symmetry of weak coherent diffraction while separating strongly asymmetric scattering contributions associated with Bragg diffraction of crystals and dynamical scattering in thick samples. Mathematical definitions and implementation details are provided in the Supplementary Materials.

### 2.8. Specimen preparation

Tobacco Mosaic Virus (TMV) samples were prepared on glow-discharged Quantifoil 1.2/1.3 grids using a Vitrobot operated at 100% humidity with 3 s blotting time. Apoferritin samples were prepared on Quantifoil 1.2/1.3 holey carbon grids and plunge frozen in liquid ethane using a Vitrobot operated at 4 °C and 100% humidity. Paclitaxel-stabilized microtubules were prepared on glow-discharged Quantifoil 1.2/1.3 grids using 5 s blotting time at 100% humidity. Collagen-mimetic peptide (CMP) nanosheets were prepared as described previously.

## 3. Results

### 3.1. Reducing beam size increases radiation tolerance

To investigate the role of beam size in radiation damage accumulation, we measured diffraction decay from protein microcrystals under parallel-beam illumination using different beam diameters. Lysosome crystals were exposed to beams with diameters of approximately 480, 160, and 80 nm, respectively. We ascertained that beam movement across the sample was minimal for these small beams, in agreement with the literature. For each condition, radial Bragg intensities from 14–20 crystals of comparable ice thickness were quantified using the reciprocal-space analysis framework described in Methods and measured as a function of accumulated dose until diffraction signal was lost (Figure 1).

**Figure 1.**
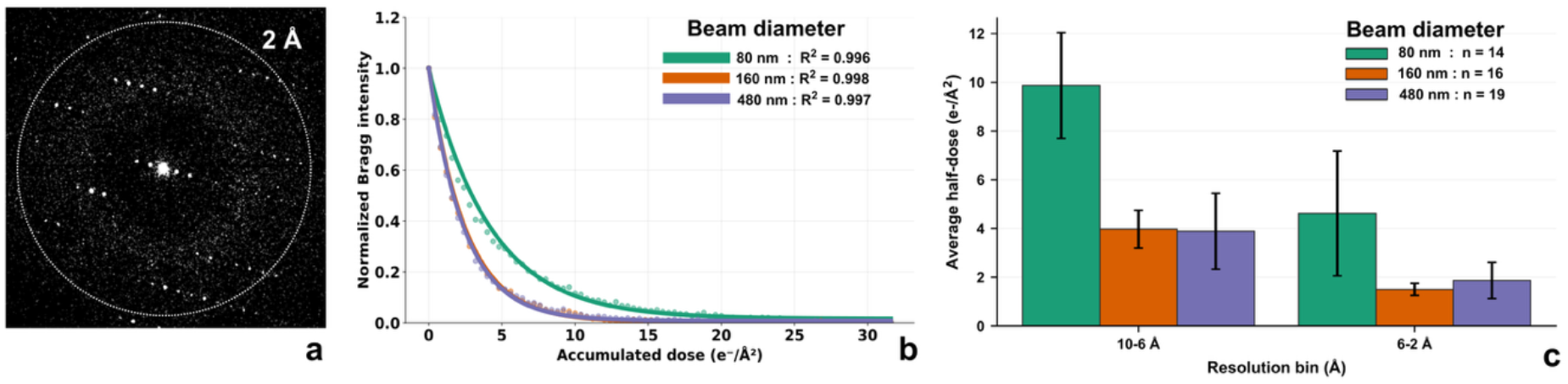
Narrow parallel-beam illumination mitigates radiation damage accumulation in protein microcrystals. (a): Representative diffraction pattern. (b): Mean normalized Bragg peak intensities as a function of accumulated dose for beam diameters of 480, 160, and 80 nm from 14–20 crystals per condition. (c): Resolution-dependent diffraction decay demonstrating preferential preservation of high-resolution information under narrow-beam illumination.

Reducing the beam diameter to 80 nm substantially prolonged the persistence of Bragg reflections compared to the larger beam conditions. In contrast, little difference was observed between the 160 and 480 nm beams. This is likely due to the fact that beam size of 480 nm exceeded the physical dimensions of the crystals in the majority of cases, resulting in effectively identical illuminated crystal volumes. These results show that radiation damage depends not only on total electron dose, but also on the spatial extent of illumination, with smaller illuminated regions preserving structural information to higher accumulated fluence.

### 3.2. Narrow parallel-beam diffraction improves reciprocal-space SNR

Previous studies using crystals reported higher reciprocal-space SNR for diffraction than for imaging [11]. To determine whether this also applies to thin non-crystalline biological specimens, we analyzed microtubule datasets acquired in both imaging and diffraction modes. Microtubules provide a particularly suitable test system because their one-dimensional translational symmetry produces discrete layer lines along the periodic axis while scattering remains continuous in the orthogonal direction. This geometry allows the reciprocal-space signal and the resolution-dependent noise and background to be estimated independently, using intensities on – and between the layer lines.

Cryo-EM imaging data acquired on a Titan Krios microscope were motion-corrected and summed to a total dose of 30 e^−^/Å^2^. The image analyzed here was recorded at relatively high defocus of about −2.4 µm to enhance contrast in the spatial frequency range considered here. Diffraction data were recorded using a 200 nm narrow parallel beam at a total dose of only 4 e^−^/Å^2^. Smaller beam diameters were not used because insufficient illuminated repeats along the microtubule axis reduced constructive interference of the layer lines. The larger beam diameter required for clear layer-line formation also increases diffuse solvent scattering and therefore favors imaging in the present comparison.

For comparison with diffraction data, cryo-EM images were Fourier transformed and converted into reciprocal-space intensity maps by squaring the Fourier amplitudes. Reciprocal-space SNR maps were subsequently calculated using the analysis framework described in Methods. SNR profiles were extracted from the microtubule equatorial layer line (Figure 2).

**Figure 2.**
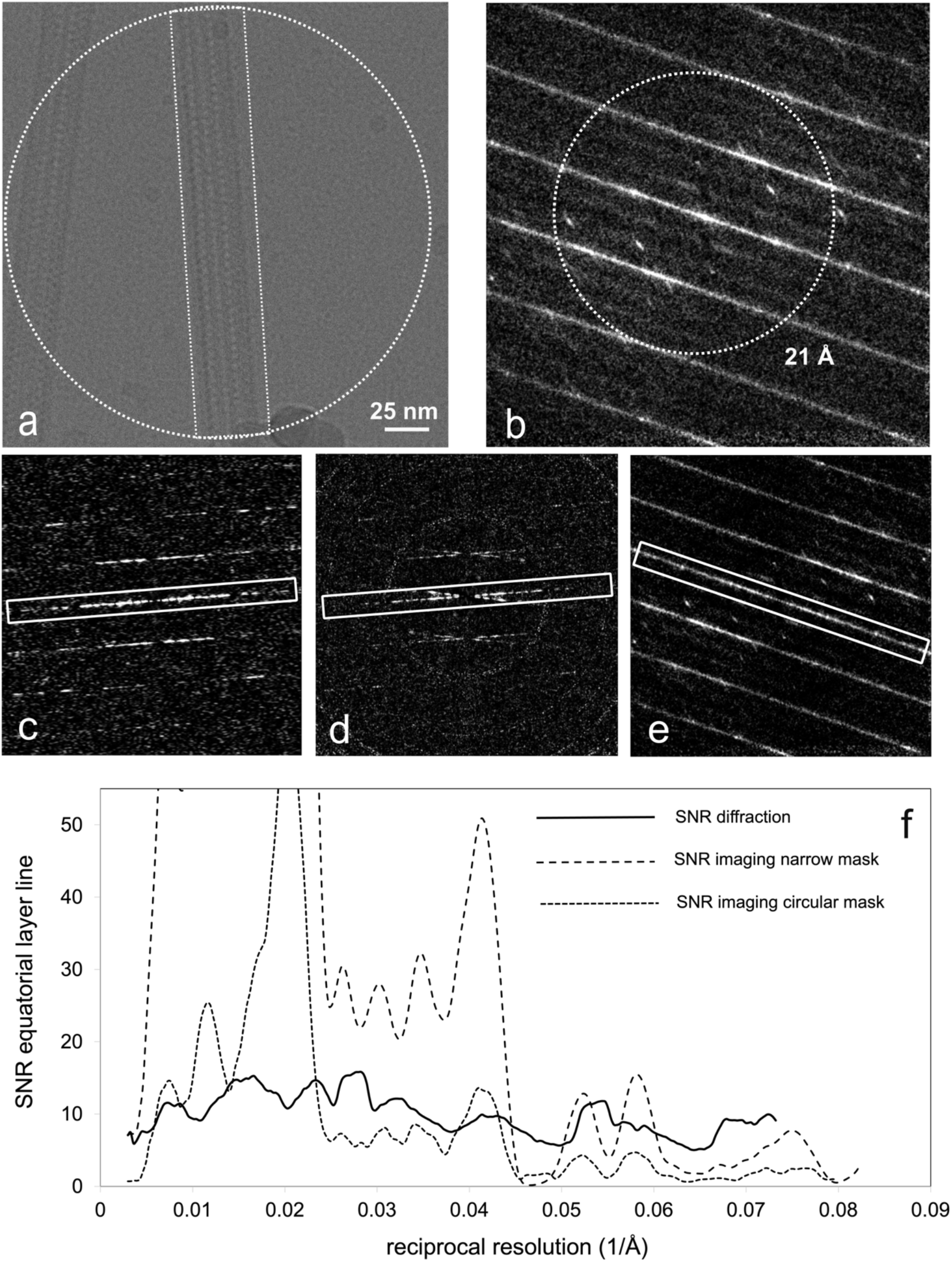
Narrow parallel-beam diffraction improves reciprocal-space SNR. (a) Motion-corrected 300 keV cryo-EM image of microtubules acquired at a total dose of 30 e^−^/Å^2^. The narrow rectangular and circular masks used before Fourier transformation are indicated. (b) Diffraction pattern of the same sample acquired using a 200 nm narrow parallel beam at 200 keV and a total dose of 4 e^−^/Å^2^. (c-e) Central sections of the reciprocal-space SNR maps calculated from the squared Fourier amplitudes of the narrow-mask image, the circular-mask image, and the diffraction pattern, respectively. All SNR maps are contoured from 0 to 5 σ. The equatorial layer lines are indicated. (f) Radial SNR profiles measured along the equatorial layer lines. Diffraction yields substantially higher SNR than imaging beyond 20 Å resolution.

Despite a 7.5 times lower dose, at resolutions higher than 20 Å, the average diffraction SNR was about five times higer than the average imaging SNR calculated using a circular 200 nm mask, and approximately 50% higher than the imaging SNR under idealized narrow-mask conditions.The circular 200 nm mask provides the most direct comparison because it matches the illuminated area used for diffraction. In contrast, the narrow rectangular mask reduces solvent contributions from the imaging data and therefore estimates the SNR improvement that could be achieved using narrower diffraction beams scanned along the microtubule axis.

### 3.3. Multimodal narrow-beam scanning enables structural analysis of biological specimens

To extend narrow parallel-beam diffraction beyond stationary measurements, we developed a multimodal scanning workflow termed 4D-para-STEM that combines diffraction and imaging acquisition from identical specimen locations under cryogenic conditions. Beam centering and parallel illumination were maintained during scans over fields of view exceeding 8 µm (Figure S2).

Initial benchmarking was performed using polycrystalline gold samples. Scan grids of 100 × 100 positions yielded 10,000 diffraction patterns that were reconstructed into vDF images by integrating annular diffraction intensities. The resulting reconstructions preserved scan geometry and allowed direct visualization of the scanned region (Figure S2).

We next applied the workflow to TMV specimens (Figure 3). Following thickness-guided selection of suitable ice regions, vDF reconstruction enabled reliable identification of scan positions containing protein signal and revealed diffraction contrast along individual TMV particles (Figure 3c).

**Figure 3.**
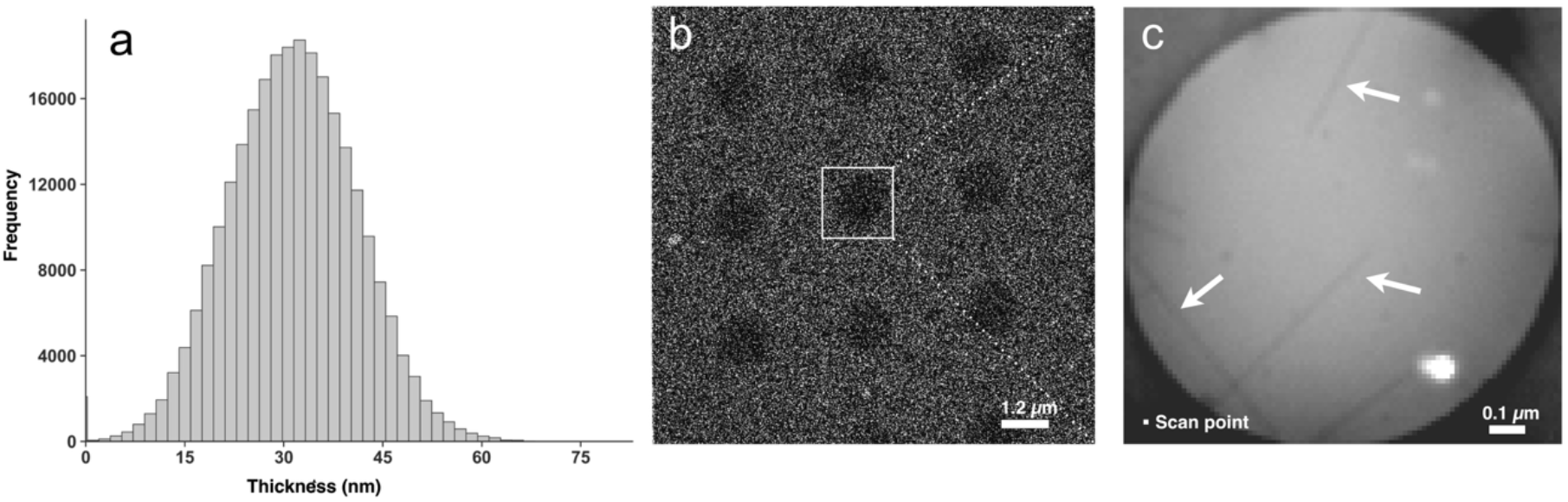
4D-para-STEM enables reliable localization of protein signal from vDF images. (a): Thickness map histogram showing the specimen thickness distribution. (b): Survey image of scan positions, providing a guideance for the subsequent scan. (c): Virtual DF detector reveals TMV positions. Arrows indicate scan points of TMVs present in the sample.

To determine whether narrow parallel-beam diffraction could recover structural information from non-crystalline biological specimens, we recorded stationary diffraction measurements from TMV-rich regions. Diffraction patterns were acquired under parallel-beam conditions from selected central specimen regions and using selected area aperture (SAA), to minimize Fresnel fringes and phase fluctuations. Individual 10 ms diffraction frames were recorded at doses of approximately 0.4 e^−^/Å^2^ per frame and summed to a total dose of 34 e^−^/Å^2^. Under these conditions, TMV diffraction features remained clearly detectable to a resolution of 4.6 Å following diffraction-frame decomposition analysis (Figure 4).

**Figure 4.**
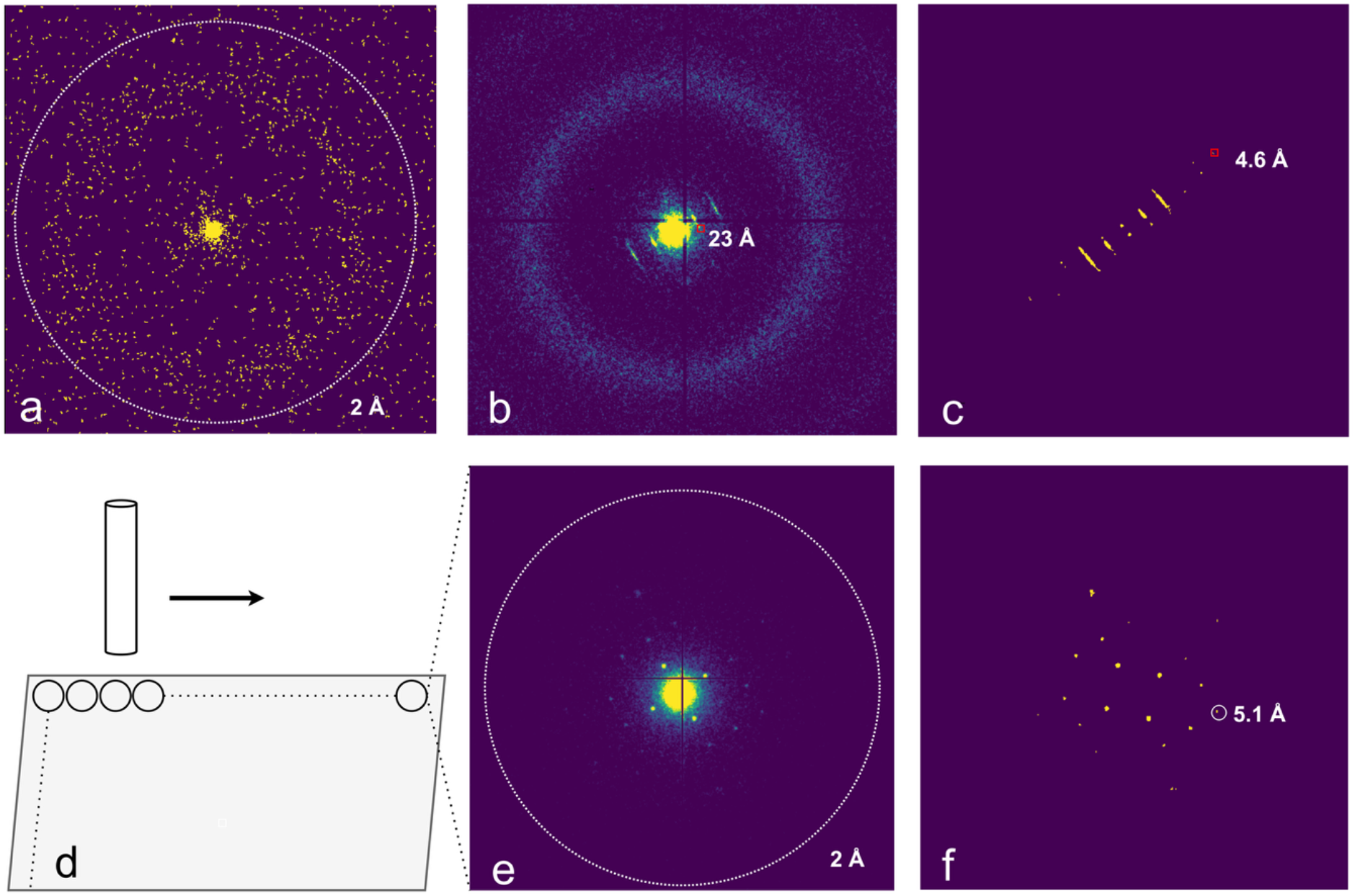
Parallel-beam diffraction recovers structural information from non-crystalline biological specimens. Top row: Analysis of diffraction patterns from selected area containing TMV. (a): An individual diffraction pattern obtained using 125nm selected area aperture and a 10 millisecond exposure time. (b): Summed diffraction pattern from TMV.(c): Decomposed frame using our decomposition approach. Bottom row : 4D-para-STEM with small parallel beam of CMPs. (d): Scan setup. (e): Individual diffraction frame. The pattern shows clearly distinguishable diffraction spacings. (f): Decomposed frame using our decomposition approach from single frame.

We next measured collagen-mimetic peptide (CMP) assemblies [12]. Raster scanning with ~40 nm beam diameters produced diffraction patterns with sharp Bragg peaks from localized regions of the peptide sheets (Figure 4). Despite accumulated doses approaching 130 e^−^/Å^2^ over multiple scans, diffraction spacings remained readily detectable, indicating that narrow parallel-beam scanning preserves interpretable structural information even after repeated scans.

Together, these results show that narrow parallel-beam illumination enables biologically relevant diffraction measurements in both stationary and scanning modes.

## 4. Discussion

Our results show that beam diameter is a major determinant of dose efficiency in cryo-EM, indicating that a narrow beam can increase the structural information obtained from a given electron dose.

Cryo-EM of biological specimens is limited by four major sources of information loss that are made worse by using a wide beam. Although these effects arise from different physical mechanisms, narrow-beam illumination reduces all four:

- *Solvent scattering*. In imaging, the defocus required for phase contrast spreads the signal on the detector, causing scattering from nearby vitreous solvent to blur into the molecular image. In diffraction, solvent background becomes even more severe. Narrow-beam diffraction avoids much of this problem because the surrounding solvent is not illuminated.
- *Beam-induced motion*. A narrow beam limits the electrostatic charging to a smaller area, thereby stabilising the specimen [8,9].
- *Radiation damage*. Although the total amount of damage is determined mainly by the number of electrons hitting the sample, part of the damage ripples out beyond the illuminated region. When a broad beam is used, much of this damage remains within the illuminated area. This is less so with a narrow beam. Similar effects were observed in STEM studies of zeolites [13].
- *Shot noise*. Radiation-sensitive samples require extreme low-dose conditions [14]. Narrow-beam illumination increases the usable dose before damage dominates, thereby improving counting statistics.

In idealized coherent-scattering theory, diffraction and imaging preserve scattered amplitudes equally well, while imaging additionally retains phase information. However, earlier studies using protein crystals showed that in practice, diffraction measurements produce substantially higher SNR [11]. Beam-induced motion, charging, contrast-transfer attenuation, coherence losses, and detector limitations substantially reduce the efficiency of real-space imaging. Our results show that the SNR advantage of diffraction is not limited to crystals, but also applies to continuous diffraction from thin, partially ordered biological assemblies such as microtubules. Furthermore, unlike images, diffraction data are constrained by Friedel symmetry, diffuse solvent isotropy, and Bragg asymmetries, which help separate structural signal from background.

Together, these results suggest that narrow-beam diffraction makes more efficient use of electrons for biomolecular structure determination. However, the signal from individual molecules remains weak, and phases are lost in diffraction. Recovering structure from single-molecule diffraction therefore requires collecting many diffraction patterns from different molecular orientations, together with methods for phase retrieval, just like cryo-EM single particle analysis.

Our multimodal 4D-para-STEM framework addresses these requirements by scanning a narrow parallel beam across the sample while collecting diffraction and imaging data from each position. We implemented Köhler illumination to produce a sharply defined illuminated region with minimal Fresnel fringes. In coherent diffraction imaging, such sharp support boundaries provide strong constraints for phase retrieval using Gerchberg–Saxton and related algorithms [15,16]. Corresponding image patches provide additional real-space phase constraints, and we previously showed that deep learning can combine diffraction and imaging data to reconstruct high-resolution structural detail [7].

4D-para-STEM is fast and fully programmable. This makes it possible to control not only the electron dose, but also the spatial distribution of exposure across the specimen. Initially, scan positions can be spaced sufficiently far apart that peripheral damage from neighbouring exposures does not overlap. Sampling density can subsequently be increased by recording between these positions, improving virtual dark-field reconstruction, and enabling stitching of corresponding image patches. The acquisition geometry itself therefore becomes part of the optimization of information transfer and radiation-damage management.

High-speed multimodal scanning with a narrow parallel beam in Köhler illumination is technically challenging. It requires precise optical scan/descan alignment, low aberration conditions and very fast detectors with high dynamic range and detective quantum efficiency. Our results show that these conditions can be achieved reproducibly under cryogenic low-dose conditions, generating large multimodal datasets. To handle the large data rates generated during acquisition, we developed TRPX [17,18], a lossless compression algorithm for high-dynamic range diffraction data and cryo-EM images that reduces data volume by up to an order of magnitude in real time. We previously published a methodology for distinguishing between diffraction data of amorphous ice, crystalline ice, and carbon [6], thus reducing the amount of data needed to be compressed and analyzed (more details are provided in the Supplementary Materials).

Together, these results show that narrow parallel-beam diffraction enables interpretable structural measurements from both crystalline and non-crystalline biological specimens under cryogenic conditions. More broadly, they show that illumination geometry strongly influences how efficiently electrons can be used for biomolecular structure determination. If combined with robust phase retrieval and high-throughput acquisition, narrow-beam diffraction could substantially improve the efficiency of cryo-EM.

## CRediT authorship contribution statement

S. Matinyan: Writing – review & editing, Writing – original draft, Software, Method development, Formal analysis, Conceptualization, Visualization, Validation. P. Filipcik: Writing – review & editing, Software, Validation. EvG: Writing – review & editing, Supervision, Method development, Data curation. J.P. Abrahams: Writing – review & editing, Visualization, Validation, Supervision, Software, Methodology, Funding acquisition, Formal analysis, Conceptualization

## 6. Supplementary materials

### 6.1. Image forming system alignment

Four distinct intermediate lens (IL) configurations were established in the image-forming system to make the detector conjugate to: (i) the diffraction plane, (ii) the image plane at 4k magnification, and (iii) the image plane at 30k and (iv) 50k magnification. Using this framework, we initially acquired diffraction patterns and subsequently applied the imaging configurations to collect real-space data from the identical scan area. Because IL configurations are mode-specific and introduce unique geometric complexities, beam deflection compensation was performed independently for imaging and diffraction scans. Notably, this acquisition sequence is modular and can be adapted or reordered to suit specific experimental requirements. Smallest parallel beam possible with this configuration and using a 10 *µ*m aperture was around 40 nm (Figure S1).

**Supplemetary Figure 1.**
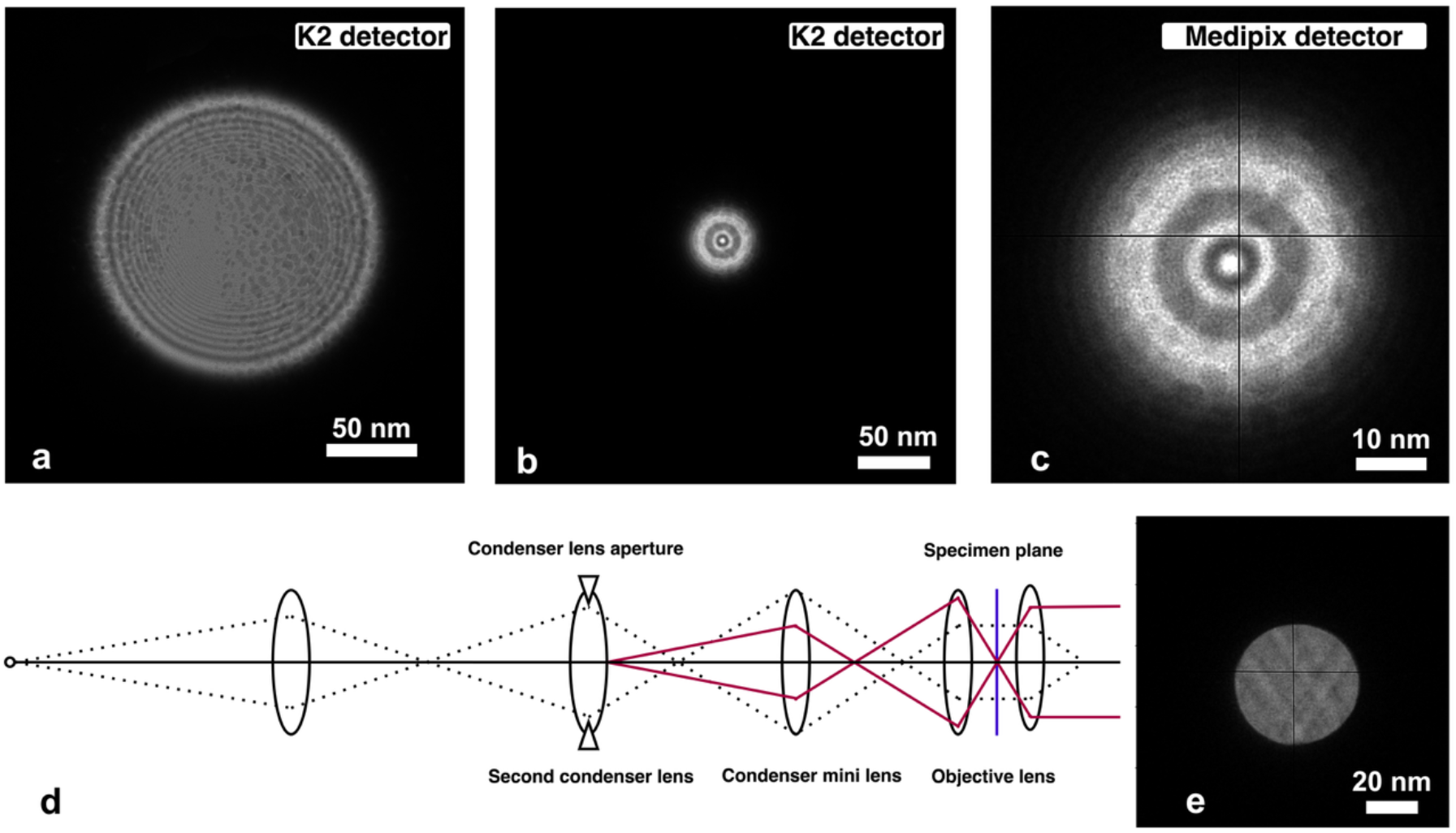
(a-c) Records of the beam on K2 and Medipix detectors, with 40 and 10 (b, c) µm CL apertures. (d) Köhler illumination. Köhler illumination is achieved by adjusting the specimen height and the excitation of the objective lens so that the CLA and the specimen are in conjugate planes. (e): Image from 4D-para-STEM scan of a microtubule specimen in Köhler illumination conditions

### 6.2. Narrow, parallel beam scan of polycrystalline samples

Scan grids of 100 × 100 positions, yielding 10,000 diffraction patterns, are shown (Figure S2). We reconstructed individual diffraction frames into virtual Dark-Dield (vDF) images by integrating annular diffraction intensities.

**Supplementary Figure 2.**
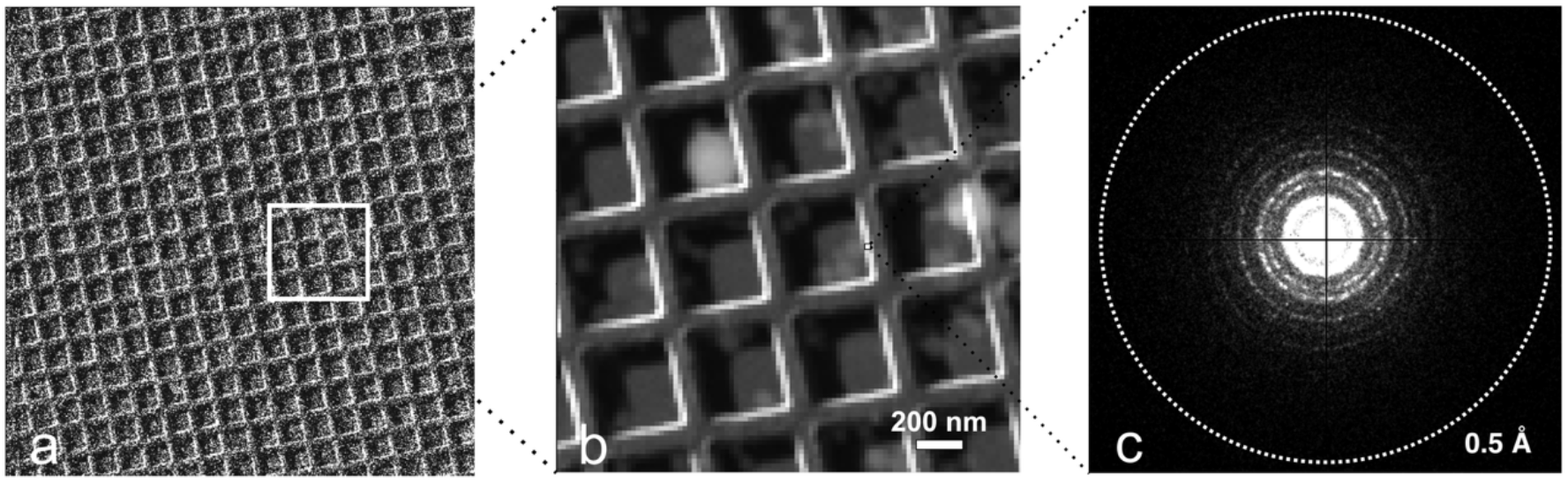
Analysis of narrow, parallel beam diffraction patterns from polycrystalline sample. (a): Field of view, from where a scan has been performed (white square). (b): A virtual DF image from a scan area, showing circular sum of diffraction intensity of each pattern as a pixel value. (c): Individual diffraction pattern.

### 6.3. Image stitching process

Each frame underwent circular masking followed by tight bounding box cropping to eliminate background pixels and preserve only the beam signal region (Figure S3). For optimal alignment, we implemented normalized cross-correlation matching combined with phase correlation for subpixel accuracy. The overlap regions were blended using gradient feathering.

The resulting stitched images enabled analysis of the complete scanned region with preserved spatial relationships.

**Supplementary Figure 3.**
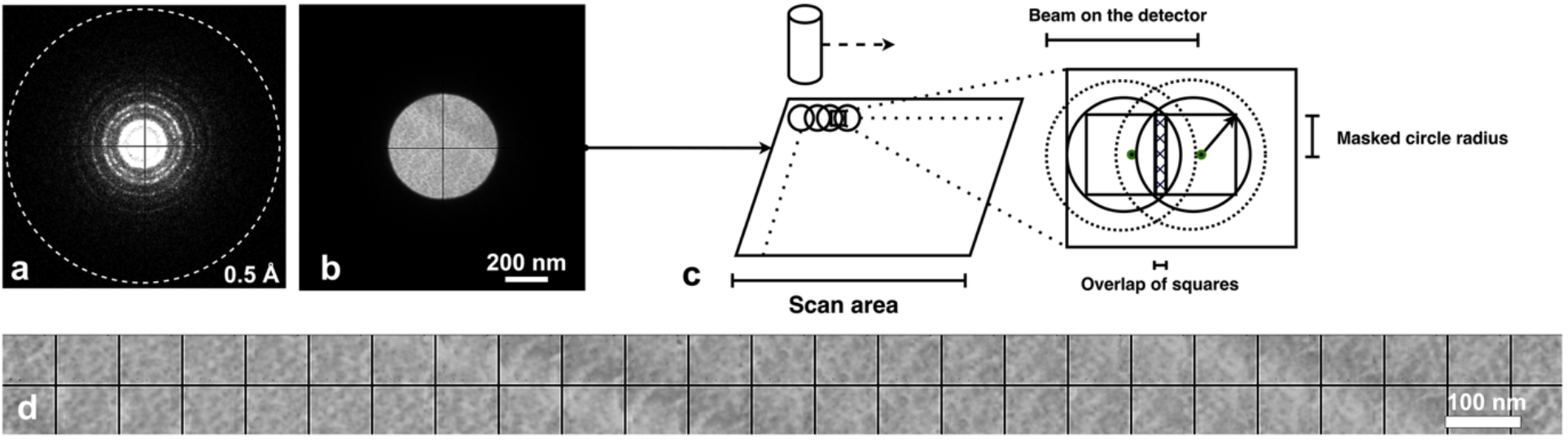
Image stitching process. (a, b) Pairs of diffraction and image scans. (c) Stitching scheme showing circular and square crops and overlap region. (d) The outcome of the stitching algorithm.

### 6.4. Image scan with Koehler illumination

Köhler illumination is an illumination technique in transmission electron microscopy (TEM) that achieves both parallel beam illumination and uniform intensity distribution across the specimen. Köhler illumination eliminates Fresnel fringes caused by the CLA. This is accomplished by carefully adjusting both the specimen position and objective lens excitation to establish a conjugate relationship between the condenser lens aperture and the specimen plane with respect to the objective lens prefield (Figure S1, bottom row). When this conjugate condition is achieved, the aperture image becomes perfectly focused at the specimen position, eliminating the non-uniform illumination and Fresnel fringe artifacts that would otherwise degrade image quality. Köhler illumination in case of narrow parallel beam becomes mandatory. In this condition image scan gives discernable details of sample positions, is not affected by Fresnel fringes, guides image analysis and provides phase information.

### 6.5. Scan and descan alignment

Descan refers to the process of realigning an electron beam that has been deflected from the optical axis using a two-stage deflector coil. This deflection occurs due to variations in the illumination position or angle of the incident electron beam on a specimen. Descan is particularly important when scanning a wide specimen area, as in the current setup, where the beam may deflect from the optical axis at the periphery of the scanned region, causing deviations in its position or angle relative to the detector. To avoid such effects, a two-stage deflector coil in the image-forming lens system is operated synchronously with the beam scan in the illumination system to always bring the electron beam back to the center of the detector.

To optimize the system for narrow parallel beam electron diffraction and imaging, we refined the descan process to minimize beam deflection from the optical axis on the detector during scanning. Because imaging and diffraction modes exhibit distinct beam paths, independent descan configurations were calibrated and stored for each modality to ensure optimal registration across all datasets.

### 6.6. Reciprocal-space signal-to-noise estimation and Bragg analysis

To quantify diffraction signal quality and radiation-induced decay, we developed a reciprocal-space analysis framework (Bragg_snr) specifically designed for weak biological nanobeam diffraction patterns, where conventional crystallographic spot-finding and indexing approaches are often unreliable (Figure S4). Standard indexing procedures generally require rotational diffraction data and globally consistent lattice geometry, whereas the present measurements consist primarily of still diffraction frames acquired from partially ordered and radiation-sensitive specimens. In addition, several biologically relevant assemblies, including helical specimens such as microtubules and TMV, produce continuous layer-line diffraction rather than discrete Bragg reflections, making conventional indexing approaches unsuitable.

The framework therefore identifies statistically significant reciprocal-space features directly from local SNR maps without requiring prior indexing or crystal orientation determination. Local reciprocal lattices are reconstructed directly from detected peaks, allowing robust identification of Bragg reflections above diffuse solvent background while accommodating multiple lattices, crystal twins, satellite lattices, partially ordered diffraction patterns, and continuous helical layer-line diffraction commonly encountered in biological specimens.

The procedure begins by transforming diffraction patterns into polar coordinates centered on the direct beam. Diffraction intensities were sampled on concentric rings spanning the selected reciprocal-space interval. At each radius, diffuse background intensity and its standard deviation were determined from the median and variance after iterative outlier rejection to suppress Bragg reflections and other localized high-intensity features. Detector gain was estimated directly from the variance-to-mean relationship of diffuse background pixels across reciprocal-space shells, assuming Poisson counting statistics. The estimated gain was subsequently used to regularize local variance estimates during reciprocal-space SNR normalization and to prevent artificial inflation of SNR values in low-count regions.

The local reciprocal-space SNR was then calculated as:

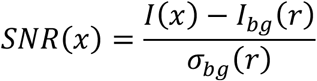

where *I*(*x*)is the measured pixel intensity, *I*_*pg*_(*r*)is the estimated radial background intensity at radius *I*_*pg*_(*r*), and *σ*_*pg*_(*r*)is the corresponding background standard deviation.

The resulting polar SNR representation was interpolated back onto the Cartesian detector grid to generate a reciprocal-space SNR map (Figure S4a, b).

Bragg reflections were identified from local maxima in the SNR map above a defined threshold within a specified reciprocal-space interval. To improve robustness against weak or partially obscured reflections, local reciprocal lattices were estimated by stacking and averaging regions surrounding strong Bragg peaks (Figure S4, c). These averaged lattice templates were subsequently used to predict additional lattice-consistent peak positions, allowing iterative expansion of Bragg masks (Figure S4, d).

Binary Bragg masks were generated from the detected reflections and used to separate Bragg and diffuse background intensities on each reciprocal-space ring. Radial Bragg intensity profiles were then calculated by integrating background-subtracted Bragg intensities as a function of scattering radius (Figure S4, e).

This framework was used for:

- reciprocal-space SNR estimation,
- diffraction decay measurements,
- radial intensity profiling,
- and comparison of imaging and diffraction signal quality.

**Supplementary Figure 4.**
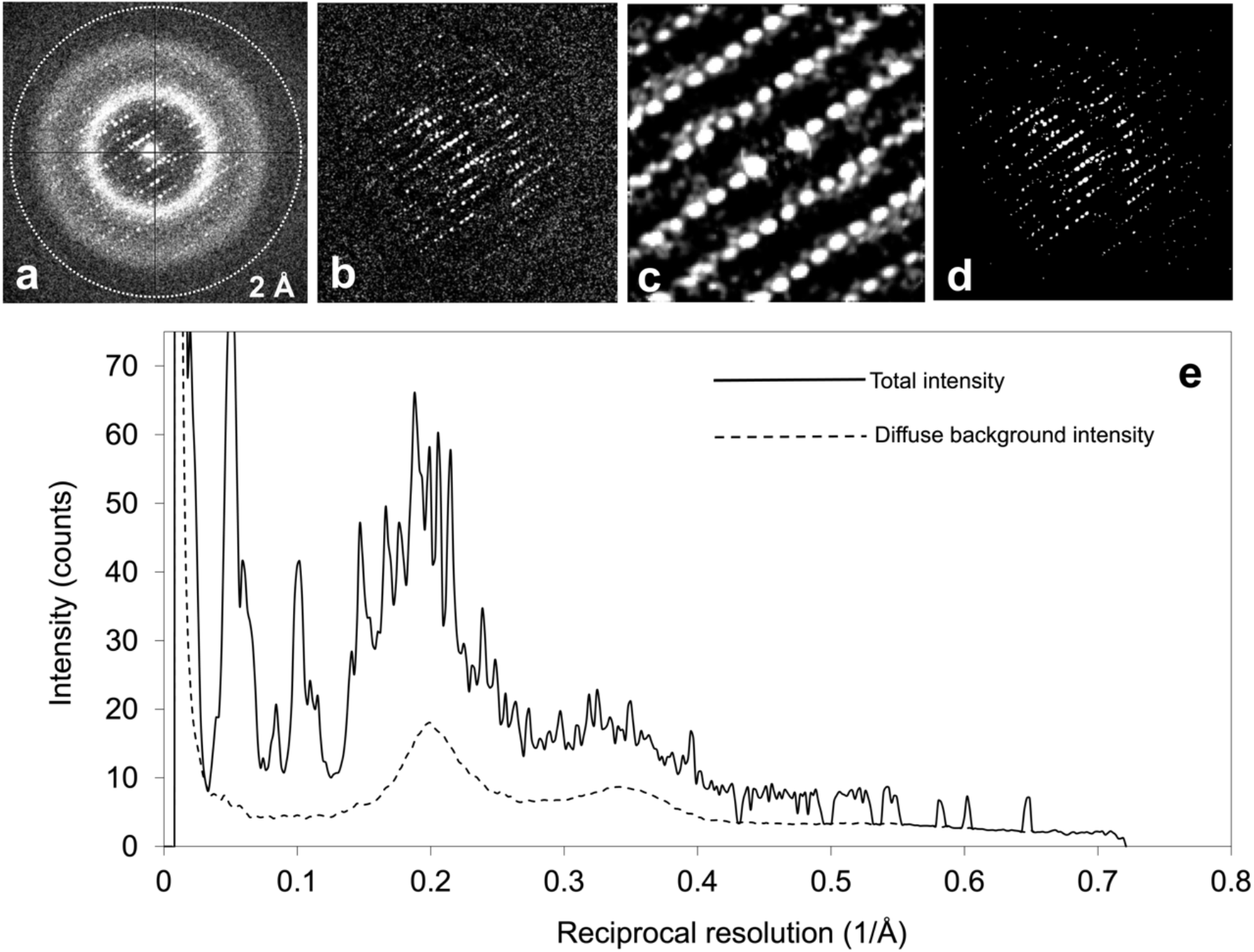
Reciprocal-space SNR analysis and Bragg peak detection workflow. (a) Raw diffraction frame from a lysosome crystal, displayed with linear intensity contours between 0 and 5× the median intensity after exclusion of the direct beam region. The diffraction pattern contains both a strong primary lattice and a weaker satellite lattice. (b) Reciprocal-space SNR map obtained after radial background estimation and variance normalization. Intensities are displayed linearly between 0 and 5*σ* above diffuse background. (c) Zoomed view of the inferred local reciprocal lattice reconstructed from high-SNR seed peaks and lattice propagation. The reconstruction identifies both the primary and weaker satellite lattice components. (d) Final Bragg mask obtained after lattice-guided peak detection and local peak expansion. Masked pixels correspond to Bragg reflections used for radial intensity analysis. (e) Radial intensity profiles showing the total integrated diffraction intensity (solid line) and the estimated diffuse background intensity (dashed line) as a function of reciprocal-space resolution. Background intensities were estimated from non-Bragg pixels following iterative outlier rejection. All contours are displayed using linear scaling within the indicated bounds.

### 6.7. Decomposition of diffraction frames of narrow beam electron diffraction

We developed a set of algorithms to analyze 4D-para-STEM data to further improve the SNR of speckle patterns. These do not in general show Bragg spots or layer lines, so cannot be analyzed using the methods in the previous section. We considered that an observed diffraction frame, 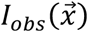, can be decomposed into four components based on specific criteria and operations. The decomposition is described as:

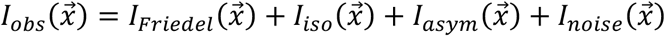

Here, the terms represent:

- 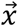 coordinate relative to the beam center
- 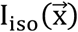 Isotropic component (radial average)
- 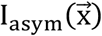 Asymmetric component
- 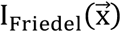 Friedel pair component
- 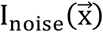 Noise component

The isotropic component primarily reflects diffuse solvent scattering and radially averaged background intensity, whereas strongly asymmetric features are typically associated with residual, contaminating Bragg reflections (for instance of crystalline water) and dynamical scattering. By contrast, non-crystalline molecular diffraction and low-resolution crystal diffraction contribute predominantly to centrosymmetric Friedel-paired intensity distributions.

The asymmetric component reflects intensities that are caused by higher resolution Bragg diffraction spots and strong dynamical scattering. They are therefore unlikely to correspond to signal of single, non-crystalline protein molecule. We consider pixel values that differ by more than a given threshold from their centrosymmetric equivalent to represent the asymmetric component. The threshold is defined by:

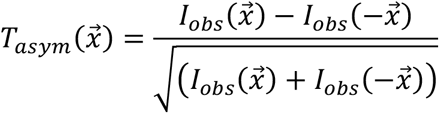

Assuming the detector gain is unity and counting statistics, the pixels for which 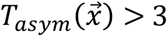 represent the asymmetric component, given by:

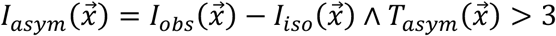

Pixels neighboring those flagged as part of the asymmetric signal are also included in the asymmetric component.

The isotropic intensity corresponds to the disordered bulk of the solvent molecules. It is the radial average of 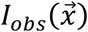 excluding regions identified as *I*_*asym*._

The Friedel pair component corresponds to the signal of the non-crystalline single molecule. It is defined by the pixels are not included in the asymmetric component and for which the intensity is of both 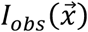 and 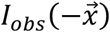 are higher than 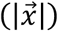:

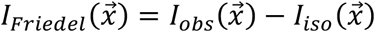

Where: 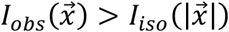 and 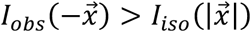 and 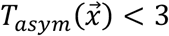.

The noise component is calculated as the residual:

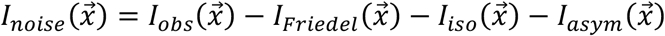

The noise component is the only component that can have both positive and negative pixel values and captures all unexplained variance. The other components are always positive. The decomposition is performed iteratively, ensuring that the components are non-overlapping and contribute additively to reconstruct 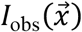. Additionally, the Friedel symmetry and radial isotropy assumptions guide the identification of *I*_asym_, *I*_iso_ and *I*_friedel_.

Applying this decomposition substantially improved the interpretability of diffraction features from both crystalline and non-crystalline biological specimens. The approach was first validated using lysosome crystal diffraction patterns and subsequently applied to TMV, and collagen-mimetic peptide assemblies (Figures 4 and Figure S5). In each case, decomposition enhanced structurally relevant diffraction features while suppressing diffuse background and non-informative scattering contributions.

**Supplemetary Figure 5.**
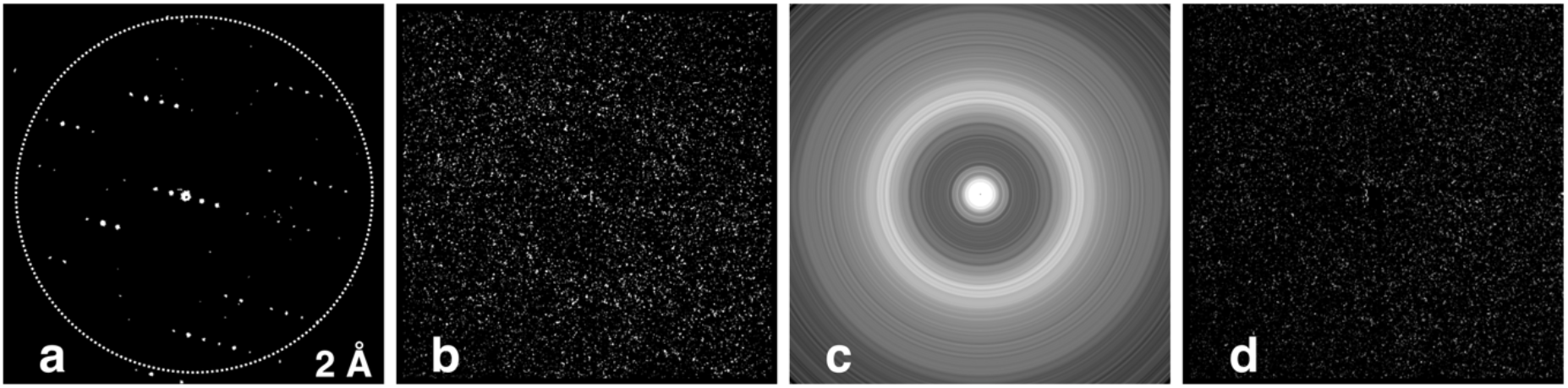
Decomposition of diffraction patterns of lysosome crystals into asymmetric (a), centrosymmetric (Friedel) (b), isotropic (c) and residual noise (d) components.

### 6.8. Machine learning based identification of diffraction scan positions containing protein signal

We previously demonstrated how we use machine learning to reduce the amount of data needed to be saved and analyzed later. Currently, the approach uses ResNet-34 architecture as the main model [20]. A graphical user interface (GUI) has been developed to allow easy labeling of scan data from vDF images (Figure S6).

To further improve our previous approach and extend to finding individual protein hits, we collected a diffraction and image scan at the same locations for apoferitin samples. Correlation analysis allowed optimizing the fit between the diffraction and imaging data. To streamline this process, a custom-designed, second GUI allowed us to overlay the image scans with calibrated scan positions (Figure S6), effectively enabling manual selection of diffraction positions corresponding to protein-containing regions.

**Supplementary Figure 6.**
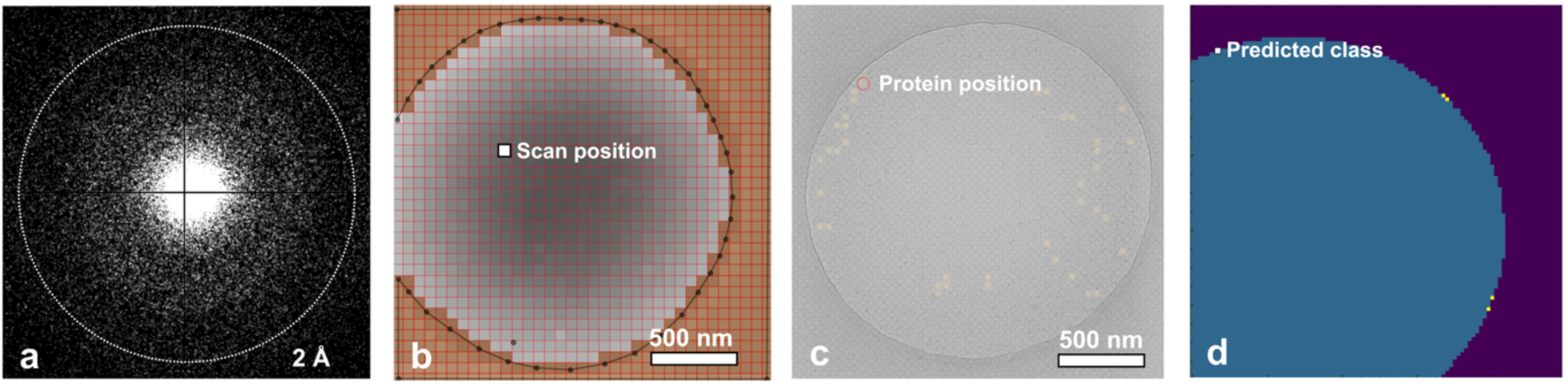
Identification of vitreous ice and protein positions from diffraction patterns and high resolution images. (a,b). The holes are identified using virtual DF and the overlaid scan positions are used to select vitreous ice and protein positions from vDF(b) and from high-resolution image data (c) respectively. Two separate GUIs were developed for this purpose (b, c). (d) Prediction results of vitreous ice positions are shown.

### 6.9. Beam and pixel size calibration

We used an imaging K2 detector in counting mode to record an image of a polycrystalline gold (Au) sample with 50k nominal magnification, 5000 *ms* exposure time and with a 40 *µm* CLA. The Fourier-transformed image showed a diffraction ring corresponding to the gold nanoparticles. We measured its diameter as a d-spacing using Adxv software [21] and calculated the magnified pixel size.

In a typical alignment, the pixel size was 0.074 nm on the K2 detector and beam diameter was 165 nm with a 40 µm CLA. We then changed the CLA to 10 µm and recorded the same beam on both K2 and Medipix detectors. In this case, we could calculate the magnified pixel size on the Medipix detector, which was 0.12 nm using these settings (Figure S1).

### 6.10. Quantification of specimen thickness

Quantifying specimen thickness is important for accurate scanning of biological samples. A common method involves analyzing Electron Energy Loss Spectroscopy (EELS) data, specifically by measuring the intensity of zero-loss electrons relative to the total electron intensity. The specimen thickness (t) in mean free paths (λ) is related to the natural logarithm of their intensity ratio. This relation provides a reliable measure of thickness based on the exponential decay of zero-loss intensity with increasing path length through the specimen. We used this approach to find suitable parts in the sample to perform the scan.

### 6.11. Source code

Source code and implementation details are available at: https://github.com/senikm/paraSTEM under MIT license.

## References

[1] T. Grant, N. Grigorieff, Measuring the optimal exposure for single particle cryo-EM using a 2.6 Å reconstruction of rotavirus VP6, Elife 4 (2015). 10.7554/ELIFE.06980.

[2] D. Shi, R. Huang, Analysis and comparison of electron radiation damage assessments in Cryo-EM by single particle analysis and micro-crystal electron diffraction, Front. Mol. Biosci. 9 (2022) 988928. 10.3389/FMOLB.2022.988928/BIBTEX.

[3] T. Bepler, K. Kelley, A.J. Noble, B. Berger, Topaz-Denoise: general deep denoising models for cryoEM and cryoET, Nat. Commun. 11 (2020) 1–12. 10.1038/S41467-020-18952-1; TECHMETA.

[4] H. Li, H. Zhang, X. Wan, Z. Yang, C. Li, J. Li, R. Han, P. Zhu, F. Zhang, Noise-Transfer2Clean: denoising cryo-EM images based on noise modeling and transfer, Bioinformatics 38 (2022) 2022–2029. 10.1093/BIOINFORMATICS/BTAC052.

[5] M.C. Kwon, J.P. Abrahams, Elastic and inelastic interactions of electrons in transmission electron microscopy, Ultramicroscopy 284 (2026) 114383. 10.1016/J.ULTRAMIC.2026.114383.

[6] S. Matinyan, B. Demir, P. Filipcik, J.P. Abrahams, E. Van Genderen, Machine learning for classifying narrow-beam electron diffraction data, Acta Crystallogr. Sect. A Found. Adv. 79 (2023) 360–368. 10.1107/S2053273323004680/HTTPS://JOURNALS.IUCR.ORG/SERVICES/RSS.HTML.

[7] S. Matinyan, P. Filipcik, E. van Genderen, J.P. Abrahams, DiffraGAN: a conditional generative adversarial network for phasing single molecule diffraction data to atomic resolution, Front. Mol. Biosci. 11 (2024) 1386963. 10.3389/FMOLB.2024.1386963/BIBTEX.

[8] P. Bullough, R. Henderson, Use of spot-scan procedure for recording low-dose micrographs of beam-sensitive specimens, Ultramicroscopy 21 (1987) 223–230. 10.1016/0304-3991(87)90147-1.

[9] K.H. Downing, Spot-Scan Imaging in Transmission Electron Microscopy, Science (80-.). 251 (1991) 53–59. 10.1126/SCIENCE.1846047;WEBSITE:WEBSITE:AAAS-SITE;JOURNAL:JOURNAL:SCIENCE;ISSUE:ISSUE:DOI.

[10] B. Küçükoğlu, I. Mohammed, R.C. Guerrero-Ferreira, S.M. Ribet, G. Varnavides, M.L. Leidl, K. Lau, S. Nazarov, A. Myasnikov, M. Kube, J. Radecke, C. Sachse, K. Müller-Caspary, C. Ophus, H. Stahlberg, Low-dose cryo-electron ptychography of proteins at sub-nanometer resolution, Nat. Commun. 15 (2024). 10.1038/s41467-024-52403-5.

[11] T. Latychevskaia, J.P. Abrahams, Inelastic scattering and solvent scattering reduce dynamical diffraction in biological crystals, Acta Crystallogr. Sect. B Struct. Sci. Cryst. Eng. Mater. 75 (2019) 523–531. 10.1107/S2052520619009661.

[12] A.D. Merg, G. Touponse, E. van Genderen, X. Zuo, A. Bazrafshan, T. Blum, S. Hughes, K. Salaita, J.P. Abrahams, V.P. Conticello, 2D Crystal Engineering of Nanosheets Assembled from Helical Peptide Building Blocks, Angew. Chemie - Int. Ed. 58 (2019) 13507–13512. 10.1002/ANIE.201906214;ISSUE:ISSUE:DOI.

[13] A. Velazco, A. Béché, D. Jannis, J. Verbeeck, Reducing electron beam damage through alternative STEM scanning strategies, Part I: Experimental findings, Ultramicroscopy 232 (2022) 113398. 10.1016/J.ULTRAMIC.2021.113398.

[14] W.T. Baxter, R.A. Grassucci, H. Gao, J. Frank, Determination of Signal-to-Noise Ratios and Spectral SNRs in Cryo-EM Low-dose Imaging of Molecules, J. Struct. Biol. 166 (2009) 126. 10.1016/J.JSB.2009.02.012.

[15] R. Gerchberg, A practical algorithm for the determination of phase from image and diffraction plane pictures, Optik (Stuttg). (1972).

[16] J.P. Abrahams, Bias Reduction in Phase Refinement by Modified Interference Functions: Introducing the γ Correction, Urn:Issn: 0907-4449 53 (1997) 371–376. 10.1107/S0907444996015272.

[17] S. Matinyan, P. Filipcik, D.G. Waterman, C.D. Owen, J.P. Abrahams, TRPXv2.0: superfast, parallel compression of diffraction patterns and images, with native Python and HDF5 support, Ultramicroscopy 281 (2026) 114298. 10.1016/J.ULTRAMIC.2025.114298.

[18] S. Matinyan, J.P. Abrahams, TERSE/PROLIX (TRPX) - a new algorithm for fast and lossless compression and decompression of diffraction and cryo-EM data., Acta Crystallogr. Sect. A, Found. Adv. 79 (2023) 536–541. 10.1107/S205327332300760X.

[19] S. Matinyan, B. Demir, P. Filipcik, J.P. Abrahams, E. Van Genderen, Machine learning for classifying narrow-beam electron diffraction data, Acta Crystallogr. Sect. A Found. Adv. 79 (2023) 360–368. 10.1107/S2053273323004680.

[20] C. Szegedy, S. Ioffe, V. Vanhoucke, A. Alemi, Inception-v4, Inception-ResNet and the Impact of Residual Connections on Learning, 31st AAAI Conf. Artif. Intell. AAAI 2017 (2016) 4278–4284. 10.48550/arXiv.1602.07261.

[21] B.T. Porebski, B.K. Ho, A.M. Buckle, Interactive visualization tools for the structural biologist, J. Appl. Crystallogr. 46 (2013) 1518–1520. 10.1107/S0021889813017858.

